# In vetro study on the toxic effects of acrylamide on blood antioxidants, ameliorative effects of vitamin C

**DOI:** 10.1101/2020.04.07.030304

**Authors:** Ahmed Ghamdi, Fahad Alenezi, Misfer Algoferi, Mohamed Alhawas, Mohamed Afifi

**Author notes:** **Corresponding author:** Mohamed Afifi, *Department of Biochemistry, Faculty of Science, University of Jeddah, Saudi Arabia.

## Abstract

**Background:** Acrylamide (ACR) is a naturally occurring, widely used compound, it is generated during cocking carbohydrate rich food at high temperature. Ingestion of large amounts of ACR underlies several health concerns and teratogenicity. Ascorbic acid (vit C) is a strong reducing agent greatly used to clean free radicals.

**Materials and methods:** Blood sample was obtained from 46 years old, healthy nonsmoking man in heparinized tubs. Blood sample was immediately divided into seven parts as triplet for each. The first one was leaved as control, 2nd, 3rd and the 4th were treated with acrylamide in a concentration of 25,50 and 100 mM respectively, the 5th, 6th and the 7th were treated with acrylamide as the mentioned concentrations and vitamin C in a concentration of 100mM. Samples (one mile litter) from each tube were taken after four and 24 hours and were used for preparation of hemolysates, that were kept at −80°C till investigation of the biochemical parameters.

**Results:** The concentrations of Malondialdehyd (MDA), nitric oxide (NO) and hydrogen peroxide (H2O2) increased in ACR and/or vit. C treated samples as compared with control. The concentration of reduced glutathione (GSH) and activities of Catalase (CAT), Superoxide dismutase (SOD), Glutathione reductase (GR), Glutathione peroxidase (GPx) and glucose 6 phosphate dehydrogenase (G6PDH) decreased significantly in ACR and/or vit. C treated samples as compared with control. Meanwhile, The concentrations of MDA, NO and H2O2 decreased in samples treated with both ACR and vit. C as compared with that treated with ACR only. The concentration of GSH and activities CAT, SOD, GR, GPx and G6PDH increased significantly in samples treated with both ACR and vit. C as compared with that treated with ACR only.

**Conclusion:** ACR produce it’s toxic effect through it’s deleterious action on the antioxidant system through induction of pro-oxidants leading to exhausting of antioxidants. Vitamin C has an ameliorative action on the deleterious action exerted by ACR through improving the balance between pro-oxidants and antioxidant.

## 1. Introduction

Acrylamide (ACR) is a chemically colorless and odorless crystalline. It is usually soluble in water and some polar solvent like ethanol, methanol and acetone. This substance is formed during the preparation of some foods at a high temperature of more than 125 degrees Celsius, by roasting for some types of foods that have a high percentage of carbohydrates such as potato chips, toast, etc. (Zamani et al. 2017). It was discovered and classified as pollutants in 2002 by the Swedish National Welfare food Committee (Fang et al. 2014). ACR can also be exposed in the environment or workplaces through air and water, and during its production or use, in addition to its presence in adhesives, cosmetics, and graphic films (Taeymans et al. 2004). Many occupational and environmental problems have been registered from the wide use of ACR which was primarily used as flocculants for clarifying drinking water (Granath et al., 2001). Indeed, ACR is used mainly in the formation of poly-acrylamides, which are widely used in plastics, paints, varnishes, adhesives and mortar. It is also applied in toiletries and cosmetics (Pingot et al., 2013). Naturally, ACR is formed through interaction of amino acids with reducing sugar. This occurs during frying, grilling, baking or roasting carbohydrate rich food as bread, potato crisps, chips crackers and french fries at temperatures above 120°c. This increased the concern about cancer risks associated with the dietary intake of fried or backed carbohydrate food (Tareke et al., 2002; Zhang et al., 2005). It was evident that exposure to large doses of ACR causes damage to male reproductive glands. Adding to this, direct ACR inhalation or skin absorption irritates the exposed tissue and can lead to nausea, sweating, speech disorders, paresthesia, numbness, myalgia, urinary incontinence and paraparesis (Alberts et al., 2002). As well as, it has a carcinogenic effect in rodents (LoPachin, 2005; Rice, 2005). Many of studies showed that ACR is absorbed rapidly and effectively by means of gastrointestinal tract (Shipp et al., 2006). It passes through the placental barrier in humans and animals, so that, maternal exposure is a relevant measure of the fetal exposure to ACR (Schettgen et al., 2004). Antioxidants may enhance or inhibit ACR, as the different antioxidants have shown different results. But if they are of an unstable type, they may turn into an oxide form and thus may lead to a reduction of ACR. A common example is vitamin C (Jin et al., 2013). Vitamin C is found in animals and plants as a naturally occurring organic compound. It works as a redox buffer that can reduce and neutralize reactive oxygen species. As a water-soluble molecule, vitamin C can function in inside and outside the cells (Bindhumol et al., 2003). Also, it is a strong reducing agent that can scavenge free radicals in different biological systems (Duarte and Lunec, 2005).This work aims to investigate the toxic effects of acrylamide on blood antioxidants and the possible ameliorative action of vitamin C.

## 2. Materials and Methods

### 2.1. Chemicals

Acrylamide and vitamin C were obtained from Sigma Aldrich (St. Louis, Missouri, United States), CAS Number: 79-06-1 and 50-81-7 respectively.

### 2.2. Sample

The study was carried out by using blood sample from healthy, 46 year old male. The criteria of acceptability to ensure reliability of the experiment were, having good health, without serious illness and receiving any medical therapy, from non-alcoholic non-smoker person.

### 2.3. Sample preparation and treatment

Blood sample was collected in heparinized tubs. Blood sample was immediately divided into seven parts as triplet for each. The first one was leaved as control, 2^nd^, 3^rd^ and the 4^th^ were treated with acrylamide in a concentration of 25,50 and 100 mM respectively, the 5^th^, 6^th^ and the 7^th^ were treated with acrylamide as the mentioned concentrations and vitamin C in a concentration of 100mM. Samples (one mile litter) from each tube were taken after four and 24 hours and were used for preparation of hemolysates, that were kept at −80C till investigation of the biochemical parameters.

### 2.4. Biochemical Investigations

Hemolysate MDA, NO, H2O2, GSH, CAT, SOD, GR, GPx and G6PDH were monitored using commercial kits (catalog numbers MD 25 29,NO 25 33,HP 25, GR 25 11, CA 25 17, SD 25 21, GR 25 23, GP 2524, PD 25 26respectively) flowing the manufacturer’s instructions. Kits were supplied by bio-diagnostic (29 Tahreer St., Dokki, Giza, Egypt).

### 2.5. Statistical analysis

All data were analyzed using a one-way analysis of variance (ANOVA) using SPSS statistical version 22 software package (SPSS, Inc, USA). The inter-grouping homogeneity was determined by Duncan’s test. Data were presented as mean ± SD and *P*<0.05 was considered statistically significant. Student *t* test was used to investigate the difference between the different periods of treatment.

## 3. Results

### 3.1. Effect of acrylamide and/or vitamin C on the hemolysate concentrations of hydrogen peroxide, malodialdehyd and nitric oxide

The hemolysate concentrations of hydrogen peroxide, Malodialdehyd and Nitric oxide increased significantly in all groups either treated with acrylamid alone or in concomitant with vitamin C as compared with control. They decreased significantly in groups treated with acrylamid in concomitant with vitamin C as compared with their corresponding ones treated with acrylamid alone. These changes were dose and time dependent (table 1).

**Table 1.**
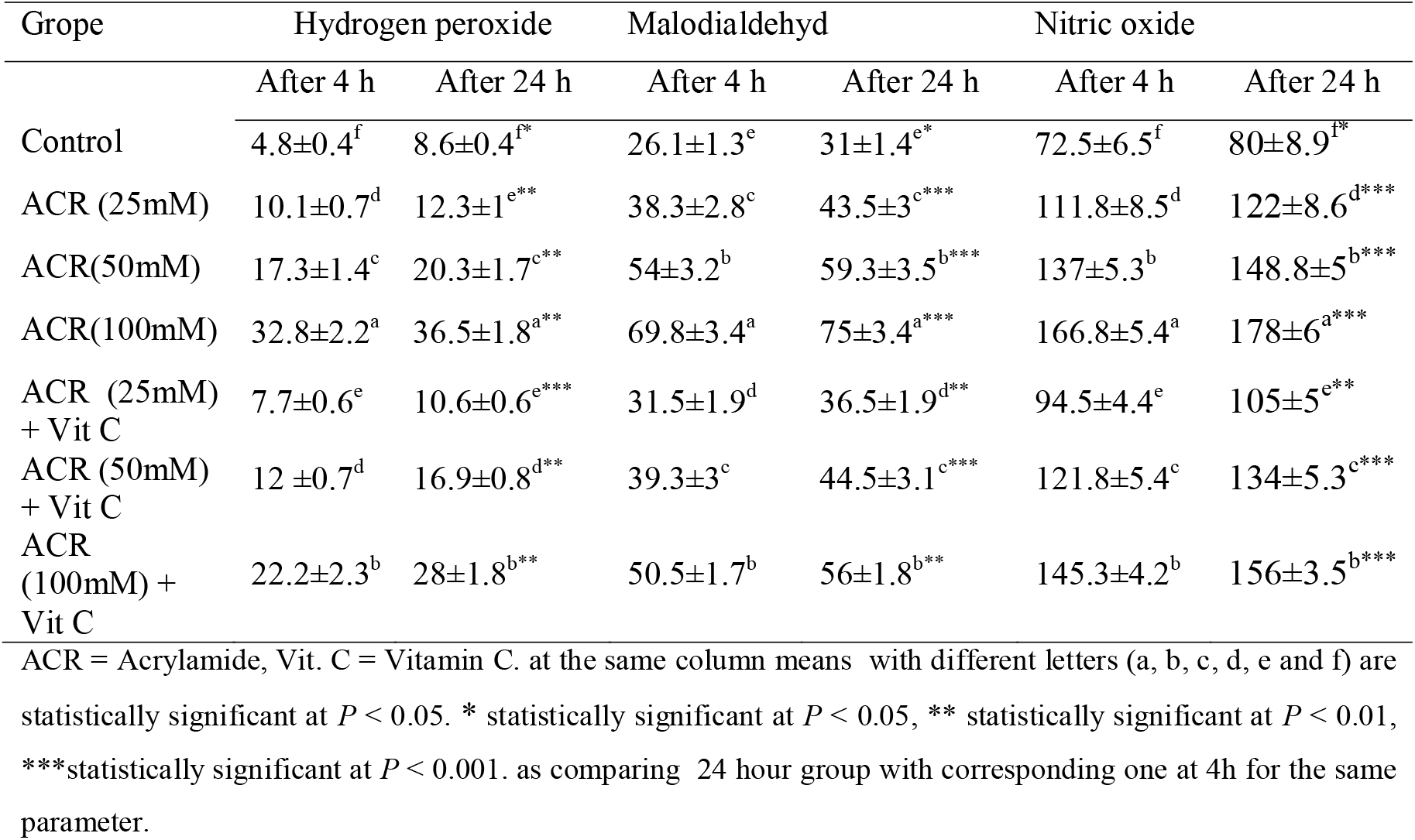
Effect of acrylamide and/or vitamin C on the hemolysate concentrations of hydrogen peroxide (uM/L) malodialdehyd (nM/dl) and nitric oxide (uM/L) after 4 and 24 hours of treatment.

| Grope | Hydrogen peroxide |  | Malodialdehyd |  | Nitric oxide |  |
| --- | --- | --- | --- | --- | --- | --- |
|  | After 4 h | After 24 h | After 4 h | After 24 h | After 4 h | After 24 h |
| Control | 4.8±0.4<sup>f</sup> | 8.6±0.4<sup>f*</sup> | 26.1±1.3<sup>e</sup> | 31±1.4<sup>e*</sup> | 72.5±6.5<sup>f</sup> | 80±8.9<sup>f*</sup> |
| ACR (25mM) | 10.1±0.7<sup>d</sup> | 12.3±1<sup>e**</sup> | 38.3±2.8<sup>c</sup> | 43.5±3<sup>c***</sup> | 111.8±8.5<sup>d</sup> | 122±8.6<sup>d***</sup> |
| ACR(50mM) | 17.3±1.4<sup>c</sup> | 20.3±1.7<sup>c**</sup> | 54±3.2<sup>b</sup> | 59.3±3.5<sup>b***</sup> | 137±5.3<sup>b</sup> | 148.8±5<sup>b***</sup> |
| ACR(100mM) | 32.8±2.2<sup>a</sup> | 36.5±1.8<sup>a**</sup> | 69.8±3.4<sup>a</sup> | 75±3.4<sup>a***</sup> | 166.8±5.4<sup>a</sup> | 178±6<sup>a***</sup> |
| ACR (25mM)<br>+ Vit C | 7.7±0.6<sup>e</sup> | 10.6±0.6<sup>e***</sup> | 31.5±1.9<sup>d</sup> | 36.5±1.9<sup>d**</sup> | 94.5±4.4<sup>e</sup> | 105±5<sup>e**</sup> |
| ACR (50mM)<br>+ Vit C | 12 ±0.7<sup>d</sup> | 16.9±0.8<sup>d**</sup> | 39.3±3<sup>c</sup> | 44.5±3.1<sup>c***</sup> | 121.8±5.4<sup>c</sup> | 134±5.3<sup>c***</sup> |
| ACR<br>(100mM) +<br>Vit C | 22.2±2.3<sup>b</sup> | 28±1.8<sup>b**</sup> | 50.5±1.7<sup>b</sup> | 56±1.8<sup>b**</sup> | 145.3±4.2<sup>b</sup> | 156±3.5<sup>b***</sup> |

ACR = Acrylamide, Vit. C = Vitamin C. at the same column means with different letters (a, b, c, d, e and f) are statistically significant at $P < 0.05$ . \* statistically significant at $P < 0.05$ , \*\* statistically significant at $P < 0.01$ , \*\*\*statistically significant at $P < 0.001$ . as comparing 24 hour group with corresponding one at 4h for the same parameter.

### 3.2. Effect of acrylamide and/or vitamin C on the hemolysate concentrations of reduced glutathione and activities of glutathione reductase and glutathione peroxidase

The hemolysate concentrations of reduced glutathione and activities of glutathione reductase and glutathione peroxidase decreased significantly in all groups either treated with acrylamid alone or in concomitant with vitamin C as compared with control. They increased significantly in groups treated with acrylamid in concomitant with vitamin C as compared with their corresponding ones treated with acrylamid alone. These changes were dose and time dependent (table 2).

**Table 2.** Effect of acrylamide and/or vitamin C on the hemolysate concentrations of reduced glutathione (mg/dl) and activities of glutathione reductase (U/dl) and glutathione peroxidase (uU/dl) after 4 and 24 hours of treatment.

| Grope | Reduced glutathione |  | Glutathione reductase |  | Glutathione peroxidase |  |
| --- | --- | --- | --- | --- | --- | --- |
|  | After 4 h | After 24 h | After 4 h | After 24 h | After 4 h | After 24 h |
| Control | 154.7±4.3<sup>a</sup> | 144.6±4<sup>a</sup> | 3.4±0.4<sup>a</sup> | 3.1±0.37<sup>a</sup> | 1027±64<sup>a</sup> | 952±38.6<sup>a</sup> |
| ACR (25mM) | 107.5±6.5<sup>c</sup> | 101.3±8<sup>c**</sup> | 2.7±0.1<sup>b</sup> | 2.4±0.17<sup>b**</sup> | 760±45<sup>c</sup> | 660±45.5<sup>c**</sup> |
| ACR(50mM) | 72.3±2.6<sup>f</sup> | 62.3±3<sup>f**</sup> | 1.9±0.1<sup>c</sup> | 1.6±0.01<sup>c**</sup> | 685±31<sup>d</sup> | 532±33<sup>d**</sup> |
| ACR(100mM) | 56.7±5.8<sup>g</sup> | 48±8<sup>g**</sup> | 1.2±0.08<sup>d</sup> | 0.9±0.01<sup>d**</sup> | 577±26<sup>e</sup> | 420±18<sup>e**</sup> |
| ACR (25mM)<br>+ Vit C | 127.3±5.3<sup>b</sup> | 117.3±5<sup>b*</sup> | 3.3±0.3<sup>a</sup> | 3±0.2<sup>a**</sup> | 952±39<sup>b</sup> | 852±38.6<sup>b*</sup> |
| ACR (50mM)<br>+ Vit C | 99.3±7.4<sup>d</sup> | 89.3±7<sup>d**</sup> | 2.6±0.2<sup>b</sup> | 2.2±0.15<sup>b**</sup> | 815±31<sup>c</sup> | 715±31<sup>c*</sup> |
| ACR<br>(100mM) +<br>Vit C | 84.3±3.5<sup>e</sup> | 74.2±3<sup>e**</sup> | 1.95±0.1<sup>c</sup> | 1.6±0.18<sup>c**</sup> | 675±49<sup>d</sup> | 575±49.3<sup>d*</sup> |

ACR = Acrylamide, Vit. C = Vitamin C. at the same column means with different letters (a, b, c, d, e and f) are statistically significant at $P < 0.05$ . \* statistically significant at $P < 0.05$ , \*\* statistically significant at $P < 0.01$ , \*\*\*statistically significant at $P < 0.001$ . as comparing 24 hour group with corresponding one at 4h for the same parameter.

### 3.3. Effect of acrylamide and/or vitamin C on the hemolysate activities of glucose 6 phosphate dehydrogenase, catalase and superoxide dismutase

The hemolysate activities of glucose 6 phosphate dehydrogenase, catalase and superoxide dismutase decreased significantly in all groups either treated with acrylamid alone or in concomitant with vitamin C as compared with control. They increased significantly in groups treated with acrylamid in concomitant with vitamin C as compared with their corresponding ones treated with acrylamid alone. These changes were dose and time dependent (table 3).

**Table 3.** Effect of acrylamide and/or vitamin C on the hemolysate activities of glucose 6 phosphate dehydrogenase (U/gHb), catalase (mU/gHb) and superoxide dismutase (U/gHb) after 4 and 24 hours of treatment.

| Groupe | G6PDH |  | Catalase |  | Superoxide dismutase |  |
| --- | --- | --- | --- | --- | --- | --- |
|  | After 4 h | After 24 h | After 4 h | After 24 h | After 4 h | After 24 h |
| Control | 13.5±1.3<sup>a</sup> | 11.6±1.7<sup>a</sup> | 307.5±17<sup>a</sup> | 277.5±15<sup>a</sup> | 32.3±4.6<sup>a</sup> | 27.3±4.6<sup>a</sup> |
| ACR (25mM) | 5.2±1<sup>d</sup> | 4.5±1.2<sup>d**</sup> | 231.7±8.9<sup>c</sup> | 201.8±8<sup>c**</sup> | 20.8±2.2<sup>c</sup> | 15.8±2.2<sup>c**</sup> |
| ACR(50mM) | 3.3±0.5<sup>e</sup> | 2.3±0.5<sup>e**</sup> | 204±7.6<sup>d</sup> | 171.5±5<sup>d***</sup> | 14.3±1<sup>d</sup> | 9.2±0.95<sup>d**</sup> |
| ACR(100mM) | 2.2±0.4<sup>f</sup> | 1.2±0.3<sup>f***</sup> | 144.6±8<sup>f</sup> | 113±8.8<sup>f***</sup> | 8.7±0.5<sup>e</sup> | 3.6±0.8<sup>e***</sup> |
| ACR (25mM)<br>+ Vit C | 8.7±0.6<sup>b</sup> | 7.7±0.5<sup>b***</sup> | 266±11<sup>b</sup> | 235±12<sup>b***</sup> | 27.7±1.8<sup>b</sup> | 22.5±1.7<sup>b***</sup> |
| ACR (50mM)<br>+ Vit C | 6.9±0.3<sup>c</sup> | 5.9±0.5<sup>c***</sup> | 230.3±9<sup>c</sup> | 201.3±8<sup>c***</sup> | 20.8±2.2<sup>c</sup> | 15.8±2.2<sup>c**</sup> |
| ACR<br>(100mM) +<br>Vit C | 4.8±0.5<sup>d</sup> | 3.8±0.6<sup>d**</sup> | 184.8±10<sup>e</sup> | 152.3±8<sup>e***</sup> | 13.5±1.7<sup>d</sup> | 8.5±1.8<sup>d*</sup> |

ACR = Acrylamide, Vit. C = Vitamin C, G6PDH = glucose 6 phosphate dehydrogenase . In the same column means with different letters (a, b, c, d, e and f) are statistically significant at $P < 0.05$ . \* statistically significant at $P < 0.05$ , \*\* statistically significant at $P < 0.01$ , \*\*\*statistically significant at $P < 0.001$ . as comparing 24 hour group with corresponding one at 4h for the same parameter

## 4. Discussion

ACR is a water soluble and easily distributed compound (Mottram et al., 2002). It is generated in starchy foods that cooked at high temperature (Erdreich et al., 2004).It is rapidly absorbed from GIT and transported to blood (Zödl et al., 2007). So blood is consider one of the main sit of ACR toxicity, there isn’t a clear mechanism that justifies ACR toxicity. We suggested that ACR produce it’s toxicity through induction the production of free radicals leading disturbance of antioxidant system. Therefore this work was designed to investigate the toxic effects of ACR on blood through monitoring the ACR effects on the prooxidants and antioxidant system of blood cells, this study is consider the first one that studied the effect of ACR on whole blood antioxidant on *invetro* level.

When there is an imbalance in the biological oxidant to antioxidant ratio, oxidative stress can occur. It could also be an initiator step to many diseases. Free radicals are constantly produced in vivo. For neutralizing of free radicals adverse effects, the body has protective barriers like antioxidant enzymes (superoxide dismutase (SOD), catalase, glutathione S-transferase, glutathione peroxidase) and proteins like GSH (Zamani et al., 2017). ACR generates free radicals disrupting the antioxidative state leading to oxidative stress (Abdel-Moneim et al., 2019). This was confirmed in the current study by increasing the blood levels of H_2_O_2_, MDA and NO, this increase is a dose and duration dependants (table 1). H_2_O_2_ is one of reactive oxygen species (ROS) that also include superoxide anion radicals (O_2_-), hydroxyl radicals (HO^·^), and singlet oxygen (^1^O_2_)are generated by oxidative stress in the body (Su-Hyeon et al., 2019). The increase of H2O2 levels in the current study indicates the ability of ACR for induction of generation of ROS that structurally damage the cell components as lipid increasing the production of MDA and proteins that manifested by the increase of blood NO levels.

The increase in free radicals and peroxide leads to induction of antioxidant system to compensate that increase till the depletion of antioxidants leading to the decrease of antioxidant levels. Blood antioxidant system include nonenzymatic antioxidant as GSH and enzymatic antoxidants as SOD, catalas, GR and GPx (Abdel-Moneim et al., 2019). Vitamin C is considered the most important water-soluble antioxidant in mammalian cells. In this study, it was confirmed that addition of vit C decrease the levels of H2O2, MDA and NO in blood samples treated with it concurrently with ACR as compared with samples treated with ACR only. Vitamin C is an effective water-soluble antioxidant found in the cytosol and extracellular fluid, is able to interact directly with free radicals, preventing oxidative damage (Rice, 2000, Banu, 2019). The content of vitamin C, a powerful antioxidant in biological fluids, functioning as a redox cofactor and catalyst, in the plasma is 20–50 μM. Ascorbate scavenge active forms of oxygen, nitrogen, and lipids such as •OH, peroxynitrite (ONOO−), LO•, LOO•, as well as antioxidant radicals such as the α-tocopheroxyl radical, thiyl radical, urate radical, beta-carotene, etc. The combination of these properties leads to the fact that vitamin C in the form of an ascorbate ion is the most important endogenous antioxidant in blood plasma that protects lipids from oxidative degradation (Linster and Van Schaftingen, 2007). Meanwhile, Vitamin C was unable to completely correct the increase of H2O2, MDA and NO as compared with control samples, this could be due to the concentration of vitamin C may be low, especially the action of vitamin C in human blood depends manly on it’s concentration, and this point need farther investigations using different doses of vitamin C.

GSH, one of the most potent reducing biological molecules, affects scavenging of free radical reactions in the erythrocytes. The consequence of the oxidative stress in cells may be the decrease of reduced glutathione and increase of its oxidized form (GSSG). Under GSH deficiency, hem iron oxidation initiates a chain reaction leading to oxidative degeneration and decomposition of hemoglobin. This process is accompanied by peroxidative damage of membrane lipids and haemolysis (Betul et al., 2009).The results showed a significant decrease in GSH in different blood samples treated with ACR in a dose and duration dependant manner (table 2). GSH is the main source of thiol groups that are required for the activity of many biological important proteins. They are also important reducing agents and cellular antioxidants. Glutathione is the principal thiol and redox buffer in mammalian cells (Tong et al., 2004). Recently it was proved that ACR is detoxicated in the body through oxidation then conjugation with GSH so it affect on GSH levels by tow ways through the direct conjugation and indirectly through generation of free radicals that neutralized with GSH all that leading to depletion of GSH (Zamani et al., 2017). GSH react with free radicals in particular H_2_O_2_ under the catalytic action of GPx leading to neutralization of free radicals to water and oxidized to oxidized glutathione (GSSG), that must reduced to GSH using the hydrogen from NADPH+H under the effect of GR (Koji and Toshio 2012).

In this study, it was confirmed that addition of vitamin C increase the levels of GSH, in blood samples treated with it concurrently with ACR as compared with samples treated with ACR only, this increase may be due to the poor ability of vitamin C for scavenger of free radicals leading to decrease of prooxidants and saving of antioxidant. Administration of vitamin C not only prevents an increase in the content of MDA in blood plasma under hypothermia of 30°C, but also contributes to a significant decrease in its level relative to the control. In addition, in RBCs, the administration of vitamin C significantly increased the GSH content in the cell during hypothermia (Klichkhanov et al., 2019). Meanwhile, Vitamin C was unable to completely correct the decrease of GSH as compared with control samples.

The author in that study proved the reduction in the activity of G6PDH in all samples treated with ACR (table 3), this is may be consider an anther mechanism for ACR action in which ACR directly inhibited G6PDH leading to a reduction of the level of reducing substance NADPH+H consequently reduced the level of GSH as mentioned before, that consider a good explanation on the depletion of GSH and increase of H2O2, MDA and NO, this result in the same line with (Birsen, 2017). G6PDH is the main source of cytoblasmic NADPH+H that necessary of GSH cycle especially in RBCs, the inhibition of G6PDH leads to reduction of NADPH+H and inhibition of GSH cycle leading to increase of free radicals and hemolysis (Hung-Chi et al., 2019). GR and GPx are important enzymes for maintaining the level of GSH, where GPx reduce the free radicals especially H2O2 in presence of GSH producing H2O and GSSG which reduced with GR to GSH. The reduction in the activities of GR and GPx in all blood samples treated with ACR in a dose and time manner in the current study (tables 2) may be due to the decrease in reducing equivalent NADPH+H that resulted from inhibition of G6PDH (Yong et al., 2009). Detoxification of H_2_O_2_ concentrations in erythrocytes is accomplished by reduction reactions catalyzed by GPx, which directs H2O2 and other peroxide attack to GSH. On the other hand, at high H_2_O_2_ concentrations, catalase plays an important role in dismutation of H_2_O_2_ to give H_2_O (Betul et al., 2009).

Among the antioxidant enzymes of erythrocytes, the most important roles in maintaining the antioxidant status are played by SOD, which provides antiradical protection and inhibits lipid peroxidation at the initiation step (Çimen, 2008). SOD is the first detoxification enzyme and most powerful antioxidant in the cell. It is an important endogenous antioxidant enzyme that acts as a component of first line defense system against reactive oxygen species (ROS). It catalyzes the dismutation of two molecules of superoxide anion (O_2_-) to hydrogen peroxide (H2O2) and molecular oxygen (O2), consequently rendering the potentially harmful superoxide anion less hazardous (Ighodaro and Akinloye 2018).. The present study reveled reduction of SOD activity in ACR treated samples (tables 3) this may be returned to exhaustion of SOD enzyme or due to damage by H_2_O_2_ that induces oxidation of amino acid residues and aggregation. Catalases catalyze direct decomposition of H_2_O_2_ to ground state O_2_. In basal conditions, catalase in erythrocytes provides protection against H2O2 generated by dismutation of superoxide radical from Hb autooxidation. The reduction in the activity of catalase in the current study in all blood samples treated with acrylamide (table 3), indicates the exhaustion of this enzymes by the over production of H_2_O_2_ ((Betul et al., 2009).

Vitamin C is considered the most important water-soluble antioxidant in mammalian cells. In this study, it was confirmed that addition of vitamin C improved the activities of all studied antioxidant enzymes, GR, GPx, SOD and catalase in blood samples treated with ACR and vitamin C as compared with that treated with ACR only. That results could explained by the ability of vitamin C to scavenger of free radical leading to correction in the disturbance between the prooxidants and antioxidants and decreasing the stress on antioxidant system. We think that, the vitamin C dose is very important and that may the cause of inability of it in our study to completely correct the disturbance caused by ACR. This point need farther investigation using different doses of vitamin C.

## Conclusion

ACR produce it’s toxic effect through it’s deleterious action on the antioxidant system through induction of pro-oxidants leading to exhausting of antioxidants. Vitamin C has an ameliorative action on the deleterious action exerted by ACR through improving the balance between pro-oxidants and antioxidant.

